# Social tension in the aftermath of public conflicts: an ethological analysis in humans

**DOI:** 10.1101/2023.10.22.563465

**Authors:** Virginia Pallante, Ivan Norscia, Marie Rosenkrantz Lindegaard

**Affiliations:** Netherlands Institute for the Study of Crime and Law Enforcement (NSCR), Amsterdam, The Netherlands; Department of Life Sciences and Systems Biology, University of Turin, Italy; Department of Sociology, University of Amsterdam, Amsterdam, The Netherlands

**Author notes:** Corresponding authors, Virginia Pallante,; De Boelelaan 1077, 1081 HV Amsterdam – The Netherlands.

**Keywords:** human behaviour, post-conflict, emotions, social tension, anger, anxiety

## Abstract

In social mammals, conflicts are stressful events for the individuals involved. In the post-conflict context, it is possible to detect the emotional state of the former opponents through the expression of displacement activities and aggressive behaviours, which indicate an increase of social tension. In humans, stressful events also induce a physiological response which leads to increased social tension behaviours. However, the variation of such behaviours in the post-conflict conflict, has never been investigated. By conducting a video analysis of street fights recorded by Close Circuit Television cameras, we assessed the variation of behaviours associated with anxiety, aggression related anger and other behaviours possibly related to both anxiety and anger (body postures and talking with gestures) in human opponents. We compared the expression of social tension behaviours before and after the eruption of the conflict and found that displacement activities (related to anxiety), aggressive behaviours (related to anger) and talking with gestures (possibly related to anxiety/anger), but not body postures, increased in post-conflict context. Moreover, displacement activities and aggressive patterns showed a temporal variation, decreasing in the 10 minutes following the conflict. Finally, different from anger-related behaviours, the level of anxiety-related behaviours was more sensitive to aggression intensity, indicating that different social tension behaviours rely on different responses that might be separated. With our study we were able to highlight the importance of the ethological approach for the study of post-conflict social tension in humans, which show a variation in its expression as observed in nonhuman primates. Following a similar comparative approach, we encourage further studies to explore the role of social tension in affecting post-conflict social dynamics.

## Introduction

In social mammals the eruption of a conflict event is followed by a period of social tension, which can reverberate to single individuals at the physiological and behavioural level (especially - although not exclusively - the opponents: Aureli, 1997; Aureli & de Waal, 2000; Aureli et al., 1989; de Waal, 2000; Judge & Mullen, 2005; Romero et al., 2009). Indeed, post-conflict social tension is detectable through the increase of individual behaviours associated with specific physiological responses (Aureli & van Schaik 1991; Aureli et al., 1989; Castles & Whiten, 1998; Cooper et al., 2007; Das et al., 1998; Fraser et al., 2008; Kutsukake & Castles, 2001; Pallante et al., 2018). One response can involve the activation of the hypothalamic–pituitary–adrenal (HPA) axis and the production of glucocorticoid hormones, which are related to stress and anxiety (a proxy for stress; Rinaldi, 2019; Sanders & Akiyama 2018; Sapolsky, 2021; Schino et al., 1996; Troisi, 2002). Another response involves the activation of the hypothalamic–pituitary–gonadal (HPG) axes and the production of testosterone, which is related with aggression and anger (Montoya et al., 2012; Peterson & Harmon-Jones, 2012). Both responses are involved in social aggression and may jointly regulate this context (Montoya et al., 2012). Consequently, it is possible for an external observer to read the emotional state of an individual after a conflict by looking at specific behavioural indicators (Troisi, 2002).

Social mammals can adopt to express distress displacement activities (Maestripieri et al., 1992), which are defined as behaviours related to anxiety – a proxy for stress involving tension and/or agitation (Barros & Tomaz, 2002; Bourin et al. 2007; Craig et al. 1995; Van Riezen & Segal, 1988) – that are displayed in a situation where they are not expected to occur (“irrelevant” to the context: McFarland, 1966; Tinbergen, 1952; Zeigler, 1964). Such activities are linked to the stress response involving the production of catecholamines and corticosteroids, together with an increased heart rate and blood pressure (Maestripieri et al., 1992). In nonhuman primates, the most common displacement activities observed during the post-conflict context are yawning, body shaking, and self-directed behaviours such as scratching and self-grooming (Castles & Whiten, 1998; Maestripieri et al., 1992; Romero et al., 2009; Troisi, 2002). Displacement activities have the adaptive function of restoring the homeostasis of individuals, ultimately leading to stress relief (Chrousos & Gold, 1992; Maestripieri et al., 1992).

Other than distress, also other basic human emotions – such as anger – are the products of evolutionarily conserved neuro-behavioural systems and can also occur in non-human mammals, where anger-like behaviours are commonly associated with aggressivity (Awathale et al., 2020; Mendl et al., 2022). In association with aggression, apes can show behavioural patterns and expressions indicative of retaliative aggressive intent which may be construed as anger (de Waal, 1988; Kret et al., 2020). Indeed, non-human primates can engage in further aggression after a conflict as an expression of social tension (Aureli & van Schaik, 1991; Castles & Whiten, 1998; Kazem & Aureli, 2005; Pallante et al., 2018; Romero et al., 2009, 2011; Schino & Marini, 2012). Such aggression can be the result of increased arousal due to a previous conflict and at the same time lead – via a negative feedback action - to stress relief (reduced cortisol level: Aureli & van Schaik, 1991; Sapolsky & Ray, 1989). As such, when opponents engage in further attacks in the post-conflict context, they show lowest rates of distress-related behaviours (Aureli & van Schaik, 1991), which is informative of the connection that exists between anxiety and anger emotional systems (Montoya et al., 2012).

Besides the mere increase of social tension behaviours, in non-human primates the expression of both anxiety-related displacement activities and anger-like manifestation (aggression) may show a decrease in the minutes following the end of the conflict (Aureli & van Schaik 1991; Castles & Whiten 1998; Cooper et al., 2007; Das et al., 1998; Koski & Sterck, 2007; Kutsukake & Castles, 2001; Pallante et al., 2018;) and a possible increase following intense aggression or other stressors (Aureli et al., 1989; Palagi & Norscia, 2011; Palagi et al., 2008; Wittig & Boesch, 2003; but see e.g., Arnold & Whiten, 2001; Wittig et al., 2015).

Similar to non-human primates, also humans show behavioural expression in relation to conflict-related anger (aggressive displays; Peterson & Harmon-Jones, 2012) and anxiety (displacement activities; Troisi, 1999). Indeed, the subjective emotional experience of anger can be linked with the changes in testosterone that occur when individuals prepare for aggression (Peterson & Harmon-Jones, 2012; Batrinos, 2012). As concerns anxiety, the first ethological studies on distress found that displacement activities are especially expressed in social tension situations in people suffering from chronic anxiety, which suggests that such activities underlie autonomic arousal (Troisi, 1999, 2002; Troisi et al., 1996). This anxiety observational assessment laid the basis for the subsequent research on the behavioural expression of distress thanks to the development of the Ethological Coding System for Interviews (ECSI), which consists in the ethogram of the displacement activities displayed by psychiatry subjects during clinical interviews. The subsequent research on the behavioural expression of distress mostly transferred the application of ECSI to other human cohorts in controlled conditions (Mohiyeddini & Semple, 2013; Pico-Alfonso et al., 2007; Sgoifo et al., 2003; Whitehouse et al., 2022; Zandara et al., 2018), but its adaptation to physiological human behaviour in natural settings has been limited so far (e.g., Barash, 1974; Shreve et al., 1988).

Ethological research on post-conflict behaviour in humans has been mostly restricted to children, who show (among others) displacement activities and anger behaviours following aggressive interactions with peers (Arsenio & Killen, 1996; Butovskaya & Kozintsev, 1999; Fujisawa et al., 2005; Ljungberg et al., 1999; Westlund et al., 2008). Children who were victims of aggression can show increased rates of displacement activities in the post-conflict context (Fujisawa et al., 2005). In humans adults, ethological research on post-conflict behaviour related to social tension is more difficult, given the rare opportunities to conduct real-life observations on naturally occurring conflict events. Two previous studies reported that people who were involved in incident events in public spaces engage in reparative strategies, showing that conflict management (i.e., consolation) and resolution (i.e., reconciliation) mechanisms follow similar dynamics to those observed in nonhuman primates (Lindegaard et al., 2017; Philpot et al., 2022). In the period following the conflict, self-oriented body postures and talking with gestures - other than displacement activities (related to anxiety) and further aggression (related to anger) - can be especially displayed by the former opponents, thus suggesting that the behavioural repertoire expressing post-conflict tension may be wider than previously described (Pallante et al., 2023). Indeed, in humans also body postures and gestures may be possibly linked to social anxiety or anger although – possibly to their variability - no clear association with a specific neuro-hormonal system has been reported (Gilboa-Schechtman & Shachar-Lavie, 2013; Lopez et al., 2017). To our knowledge, in humans there is no study on the effect of conflict intensity or clearly showing a temporal variation of social tension behaviours after a conflict, although it is known that individual homeostasis is resumed within minutes (due to the inhibition of the physiological responses related to anxiety and anger: Herman et al., 2016; Marques et al., 2022).

Despite the previous studies reporting on the link between certain behavioural patterns and social tension in humans (e.g. Troisi, 2002), thus far no quantitative analysis to compare their occurrence in the post-conflict period (compared to a baseline condition) has been run. In contrast, in non-human primates (spanning strepsirrhines and haplorrhines; e.g., brown lemurs, *Eulemur rufus x collaris*: Palagi & Norscia, 2011; Tonkean macaques, *Macaca tonkeana*: Palagi et al., 2014; chimpanzees, *Pan troglodytes*; Fraser et al., 2008), the Post-Conflict (PC)/Matched-Control (MC) method has been successfully applied to evaluate whether behaviours related social tension would increase in the minutes following a conflict (PC) compared to the baseline (MC). This method had been originally developed to compare the occurrence of affiliative behaviours observed in the post-conflict period with baseline data in order to detect possible conflict resolution mechanisms (e.g., reconciliation; de Waal & Yoshihara, 1983).

By applying the PC-MC method on adult humans, in this study we aim at assessing: i) the possible variation of social tension related behaviours around a conflict; ii) the factors possibly modulating such variation, when present. Based on the previous framework we formulated the following predictions.

### Prediction 1 - Social tension behaviours increase after conflict

Social tension in non-human primates can lead to an increase in anxiety and aggression behaviours - also as an anger-like expression - and in humans can be associated with anxiety and anger related behaviours (e.g., Aureli & van Schaik, 1991; Castles & Whiten, 1998; Kazem & Aureli, 2005; Kret et al., 2020; Montoya et al., 2012; Pallante et al., 2018; Romero et al., 2009, 2011; Schino & Marini, 2012). Thus, we expect that in humans– in the tension period following a conflict – there may be an increase in the expression of behaviours associated with anxiety (Prediction 1a), aggression related anger (Prediction 1b) and other behaviours possibly related to both anxiety and anger (self-directed body postures and talking with gestures; Prediction 1c).

### Prediction 2 - Temporal variation of post-conflict social tension behaviours

In nonhuman primates, both displacement activities and aggressive levels in the opponents decrease over time as the minutes following the end of the conflict elapse (Aureli & van Schaik 1991; Castles & Whiten 1998; Cooper et al., 2007; Das et al., 1998; Koski & Sterck, 2007; Kutsukake & Castles, 2001; Pallante et al., 2018). In humans – as in other mammals - there is a physiological inhibition – within minutes - of the hormonal cascades that generate anxiety (linked to distress) and anger related responses (Herman et al., 2016; Marques et al., 2022). Hence, we predict a decrease after a conflict - over time – of anxiety behaviours (Prediction 2a), aggression related anger (Prediction 2b) and self-directed body postures and talking with gestures (Prediction 2c).

### Prediction 3 – Effect of aggression intensity on post-conflict social tension behaviours

Non-human primates can modulate the expression of the post-conflict tension behaviours depending on the intensity of the previous conflict. Although the difference is not always obvious (possibly due to low behavioural rates, small sample size or intensity categorization; e.g. see Arnold & Whiten, 2001; Kutsukake & Castle, 2001; Wittig et al. 2015), there is evidence that in non-human primates anxiety related behaviours may increase in case of intense conflicts or stressors (Aureli et al., 1989; Palagi & Norscia, 2011). Moreover, opponents can be more likely to redirect their aggressive motivation (possibly linked to anger, de Waal 1988; Kret et al., 2020) against other group members after being involved in a severe rather than in a low intensity aggressive interaction (Palagi et al., 2008; Wittig & Boesch, 2003). In certain cohorts of humans, anxiety particularly increases after physical aggression (compared to non-physical one leading to other emotional states; Bernaldo-De-Quirós et al., 2015). Hence, we predicted an increase in anxiety-related behaviours (Prediction 3a), anger-related behaviours (Prediction 3b) and other behaviours (postures and talking with gestures, possibly linked to anxiety or anger; Gilboa-Schechtman & Shachar-Lavie, 2013; Lopez et al., 2017; Prediction 3c).

## Methods

### Video sample and subjects

We analysed CCTV footage of inter-personal conflicts in public recorded with cameras owned by the City of Amsterdam (the Netherlands), and stored by the Amsterdam police from March to June 2020. We were granted permission to use the footage for scientific purposes serving the public good, conditioned by strict security measures, as defined by the Dutch Prosecution Services (PaG/BJZ/49986). The study was postively assessed by the Ethics Review Board of the Faculty of Social and Behavioral Science at the University of Amsterdam (2022-AISSR-15534).

We instructed camera operators employed by the City of Amsterdam to notice down moments of conflicts defined as simple quarrels indicated by aggressive gesturing and frustration to serious violence involving hitting and licking someone on the ground. For each conflict, 30 minutes before and after the event were recorded, so that each clip had a duration of approximately one hour. The footage did not include sound.

From a total sample of 500 recorded incidences, we selected those which met the following criteria:

1. The footage were of a quality (resolution and brightness) high enough to allow for the coding;
2. At least one opponent was present and clearly visible for at least one minute before and one minute after the end of the conflict.

The total sample consisted in 24 clips which involved 36 opponents, including 2 people that we identified as women, and 34 as men, as assessed based on visual cues. The conflicts were recorded by public surveillance cameras that were located in different neighbourhoods of Amsterdam, including the inner entertainment areas, central business districts and suburban locations. Cameras were placed in public streets, parks, squares, pedestrian walkways, transport station exteriors and outdoors spaces of shopfronts and drinking venues. The conflicts recorded consisted mainly in street fights or disputes that resulted from traffic accidents and quarrels occurring outside locals. Conflicts occurred both during the day and in the night-life settings. Because the footage recorded covered a time span of several months that were characterized by periods of lockdown due to the Covid19 pandemic, some conflicts occurred in presence of several people in the street, while for some others bystanders were virtually absent.

### Operational Definition and Coding procedure

We constructed a data set by first identifying the conflict and defining its intensity. A conflict was identified as such when one person show at least one aggressive behaviour directed against another. A conflict was defined of low intensity when the two opponents engaged in a quarrel that did not involve any physical contact but was characterized by the display of aggressive gestures (Table 1) and invading the other’s space (within 1m). In a medium intensity conflict the opponents performed aggressive behaviours that included a physical contact such as slapping, pushing, pulling, kicking, and hitting (Table 2). High intensity conflicts involved repeated (more than one) contact patterns with one opponent directing multiple aggressive acts toward another opponent who did not fight back and showed submissive behaviours (e.g. laid on their knees or on the ground, Figure 1).

**Table 1.** The ethogram used for coding distress related behaviours, grouped per category (for the definition of the behaviours see Pallante et al., 2023).

| Behavioural categories | Behaviours included |
| --- | --- |
| Category 1. Anxiety-related behaviours (displacement activities) | Fixing hairs, Fumble, Holding the body, Moving on the place, Pacing, Redirecting attention, Self-touching, Smoking, Touching objects |
| Category 2. Anger-related behaviours (aggressive patterns) | Aggressive redirecting attention, Talking with aggressive gestures (Pointing against someone, Swinging arms, Spreading arms), Walking angry |
| Category 3. Other behaviours (postures and talking with gestures) | Postures: Arms crossed, Hands in pockets, Hands on hips, Leaning<br>Gestures employed while talking: Opening the arms, Pointing |

**Table 2.** Levels of conflict intensity.

| Level of severity | Behaviours included |
| --- | --- |
| Level 1 - low | Invading space, displaying aggressive gestures (e.g., pointing at person, feint, obscene gestures), aggressive verbal interaction including shouting and yelling, and any kind of non-physical aggressive action |
| Level 2 - mild | Isolated kicks, punches, slaps, shoves, head-butts, aggressive pushes, aggressive pulling, and any kind of aggressive action involving physical contact not protracted in time |
| Level 3 - high | Repetitive kicks, punches, slaps, shoves, head-butts, flurries of hits, aggressive pushes, aggressive pulling and any kind of prolonged aggressive action involving physical contact. |

**Figure 1.**
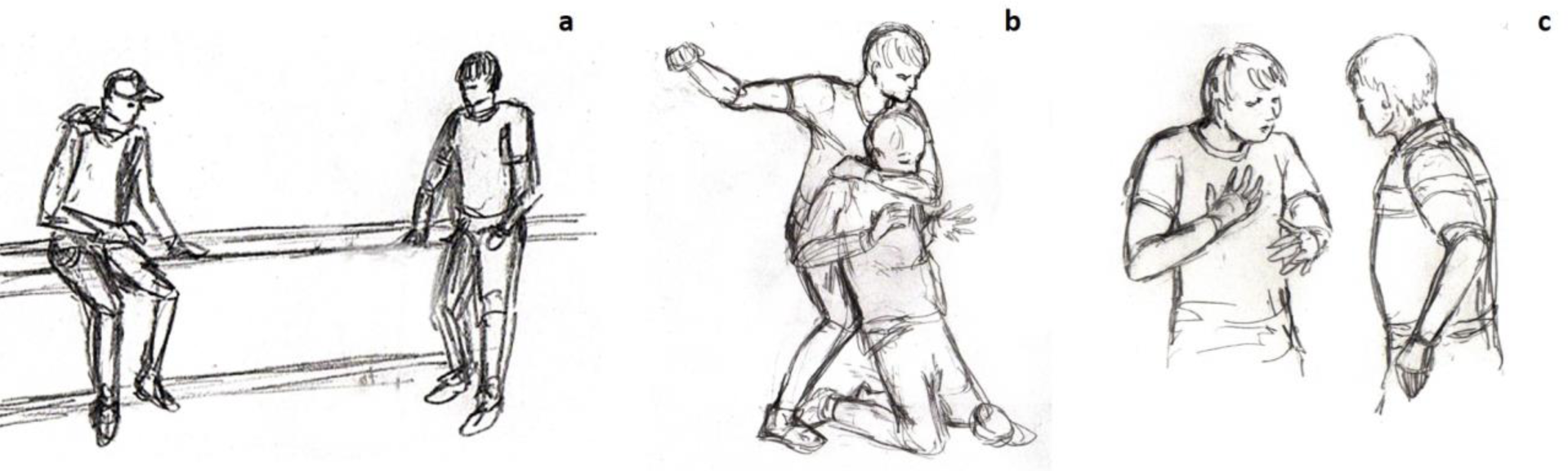
Opponents before (a) and during (b) the conflict. In c, one of the two opponents is interacting (talking with gestures) with a police officer in the post-conflict context.

Our second step in constructing a dataset was to focus on behaviour of the former opponents in the aftermath of the conflict. Opponents were defined as the individuals who engaged in the previous fight. We restricted our observations to the before-conflict (matched-control, MC) and post-conflict (PC) contexts. The MC context was defined as the period that immediately preceded the eruption of the conflict. We considered valid MC contexts only the periods where the opponents did not engage in any form of aggressive encounter. The MC context ended as soon as one of the opponents started to aggressively interact against the other. The PC context started as soon as the conflict was over, i.e., when the opponents stopped to aggressively interact and moved their attention away from each other (i.e., orienting the body and gaze away from the previous opponent). We conducted our observations on one opponent per time for at least one minute in the PC and one minute in the MC. All the behaviours displayed by the opponents in the PC and in the MC were coded according to the ethogram of the behaviours indicating a state of social tension (Table 1). We checked the reliability with a second independent coder by double-coding and comparing 10% of the videos via Cohen’s Kappa (κ), and we obtained an interobserver reliability score of 0.76 (substantial agreement).

### Statistical analysis

We conducted two rows of Generalized Linear Mixed Models (GLMMs). With the first row of GLMMs we checked whether the conflict affected the probability of observing the behaviour of interest by comparing the PC/MC contexts. In particular, models were run for anxiety related behaviours (GLMM_1_), anger behaviours (GLMM_2_), postures (GLMM_3_), and talking with gestures (GLMM_4_). In all models, we entered the occurrence of the behaviour of interest as target, binary variable (absence/presence; N=362 records) and the context as the fixed factor (binary, PC/MC). For the behaviours on which the context had a significant main effect, we ran a second raw of GLMMs to check whether – in PC - timing (minute), conflict intensity influenced the probability of observing the behaviour of interest. In particular, we ran models for anxiety related behaviours (GLMM_5_), anger behaviours (GLMM_6_), and talking with gestures (GLMM_7_). In these models we entered as dependent, binary variable the occurrence of the behaviour in the PC (absence=0; presence=1; N=226 records). Moreover, we included as fixed factors the minute of observation (numeric variable, from 1 to 10) and the intensity of the conflict (multinomial; low/medium/high intensity). We included sex as control fixed factor. For all models, the identity of the opponents was entered as a random factor.

We run the analysis with R 4.1.3 version. The models were fitted in R [(R Core team, 2019); version 3.5.3] by using the function glmer of the R-package lme4 (Bates et al., 2015). As a first step we verified if the full model significantly differed from the null model, including only the random factors (Forstmeier & Schielzeth, 2011). The likelihood ratio test (Dobson, 2002) was used to test this significance (ANOVA with argument “Chisq”). Subsequently, by using the R-function “drop1,” the p-values for the individual predictors based on likelihood ratio tests between the full and the null model were calculated (Barr et al., 2013). As the response variable was binary, a binomial error distribution was used (link function: logit). A multiple contrast package (multcomp) was used to perform all pairwise comparisons for each involvement category of significant fixed factors with the Tukey test (Bretz et al., 2010). The Bonferroni adjusted p-values were reported, along with estimate (Est), standard error (S.E.), and z-values.

## Results

GLMM_1_ (anxiety behaviour), GLMM_2_ (anger behaviour), and GLMM_4_ (talking with gestures) the full model including the fixed factor (PC/MC context) significantly differed from the null model including the random factor only (GLMM_1_:likelihood ratio test: χ^2^=4.417, df=1, p = 0.036; GLMM_2_:likelihood ratio test: χ^2^=26.215, df=1, p<0.001; GLMM_4_:likelihood ratio test: χ^2^=6.467, df=1, p = 0.011; Figure 2). Hence, the variance explained by the test predictors as a collective was significantly different from the variance explained by the variables in the null model. Therefore, we proceeded with the drop 1 procedure to check for the effect of the individual predictor. We found that the context had a significant effect on the presence of the anxiety behaviour (GLMM_1_: p=0.035; Table 3; Figure 2a), anger behaviour (GLMM_2_: p=0.002; Table 3; Figure 2b), and talking with gestures (GLMM_4_: p=0.014; Table 3; Figure 2c), which were more frequent after than before the conflict. Hence, the conflict appeared to induce the above behaviours in the subjects included in this study. For GLMM_3_ (postures) we found no significant difference between the full and the null model (likelihood ratio test: χ^2^=0.001, df=1, p=0.987; Table 3; Figure 2d) so the context (PC/MC) had not effect on the posture presence and we did not proceed with the drop 1 procedure.

**Figure 2.**
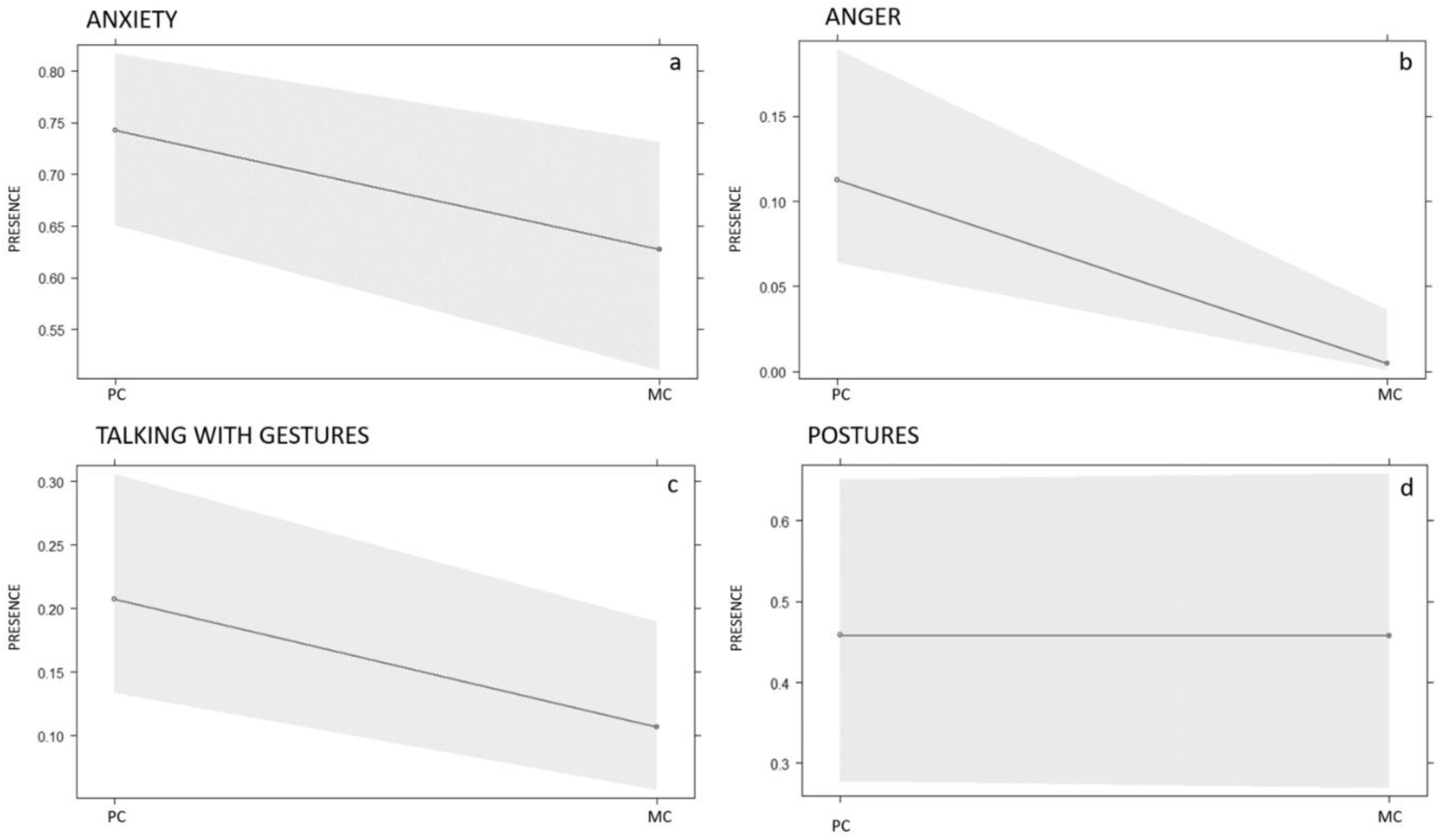
Variation of social tension behaviours in the Post-conflict (PC) compared to Matched-control (MC) context.

**Table 3.** Full results of: the influence of the context (PC/MC) on the presence of anxiety behaviour (GLMM_1_); anger behaviour (GLMM_2_); postures (GLMM_3_); talking with gestures (GLMM_4_). For all models the opponent’s identity was included as random factor. The sample size is of N=362 records for all models.

| Predictors | Estimates | SEM | CI<sub>95</sub> | Effect size | $\chi^2$ | p |
| --- | --- | --- | --- | --- | --- | --- |
| **GLMM<sub>1</sub>** | Full vs. null model: $\chi^2=4.417$, df=1, p = 0.036 | | | | | |
| (Intercept) | 1.060 | 0.224 | 0.62, 1.50 | 1.06 | 4.735 | **0.000** |
| Condition (PC/MC) | -0.539 | 0.256 | -1.04, 0.04 | -0.54 | -2.105 | **0.035** |
| **GLMM<sub>2</sub>** | Full vs. null model: $\chi^2=26.215$, df=1, p<0.001 | | | | | |
| (Intercept) | -2.065 | 0.313 | -2.68, -1.45 | -2.07 | -6.609 | **0.000** |
| Condition (PC/MC) | -3.301 | 1.041 | -5.34, -1.26 | -3.30 | -3.171 | **0.002** |
| **GLMM<sub>3</sub>** | Full vs. null model: $\chi^2=0.001$, df=1, p=0.987 | | | | | |
| (Intercept) | -0.165 | 0.402 | -0.95, 0.62 | -0.17 | -0.411 | 0.681 |
| Condition (PC/MC) | -0.004 | 0.281 | -0.55, 0.55 | -4.37e-03 | -0.016 | 0.988 |
| **GLMM<sub>4</sub>** | Full vs. null model: $\chi^2=6.47$, df=1, p=0.011 | | | | | |
| (Intercept) | -1.343 | 0.268 | -1.87, -0.82 | -1.34 | -5.014 | **0.000** |
| Condition (PC/MC) | -0.785 | 0.318 | -1.41, -0.16 | -0.78 | -2.466 | **0.014** |

For GLMM_5_ (target variable: presence/absence anxiety behaviour) the full model (including all fixed factors: timing, opponents’ sex, conflict intensity) significantly differed from the null model including the random factor only (likelihood ratio test: χ^2^=11.885, df=4, p<0.05). Timing (minute) and conflict intensity had a significant effect (p<0.05) on the anxiety behaviour presence whereas the opponents’ sex had no significant effect (see full results in Table 4). In particular, anxiety behaviour were lower for medium intensity conflicts (Figure 3; Tukey test: medium-low: Est=-0.999; S.E.=0.455, z=-2.197, p=0.071; high-low: Est=0.414; S.E.=0.432, z=0.957, p=0.603; high-medium: Est=1.412; S.E.=0.483, z=2.924, p=0.010) and decreased as the time from the conflict elapsed (Figure 4).

**Figure 3.**
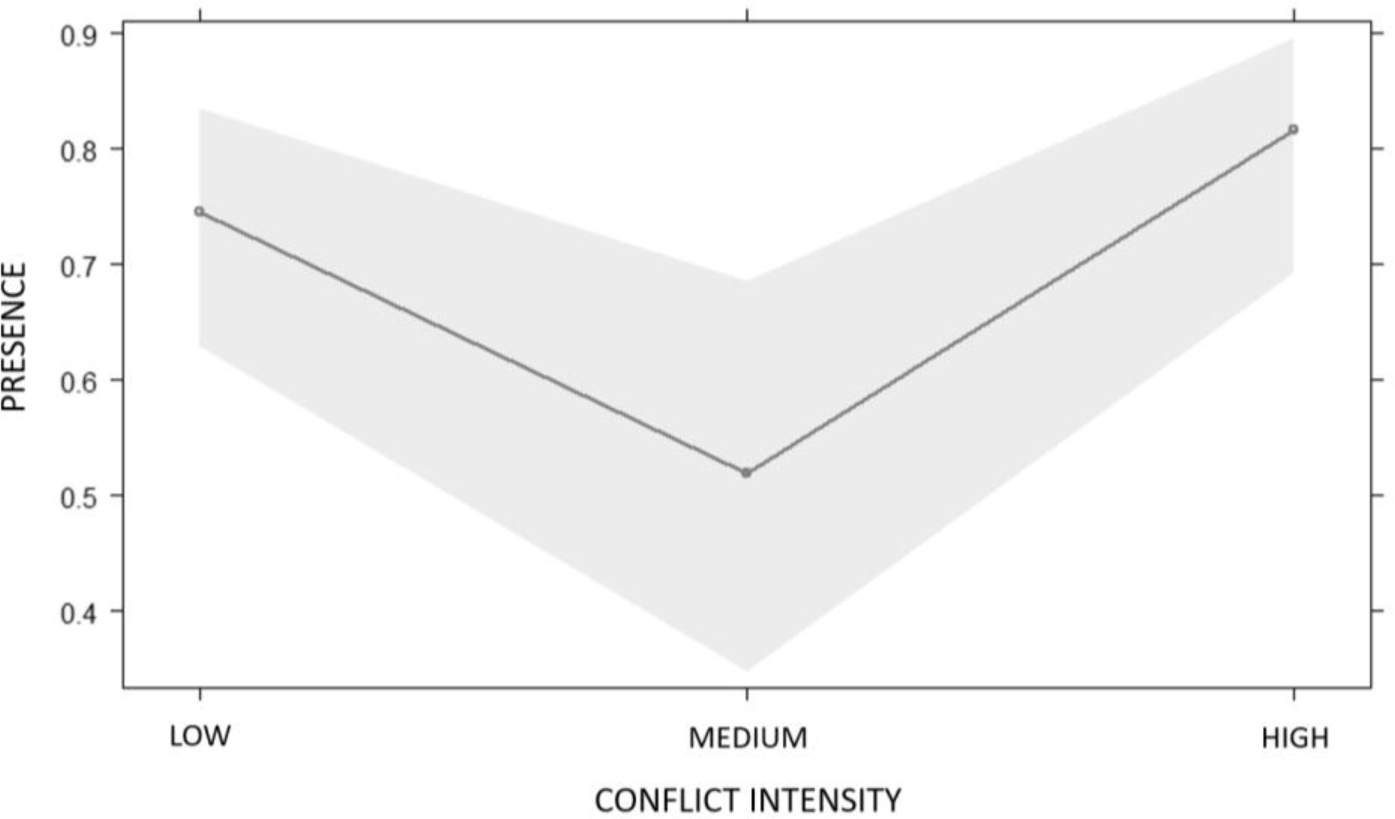
Effect of intensity on anxiety-related behaviours.

**Figure 4.**
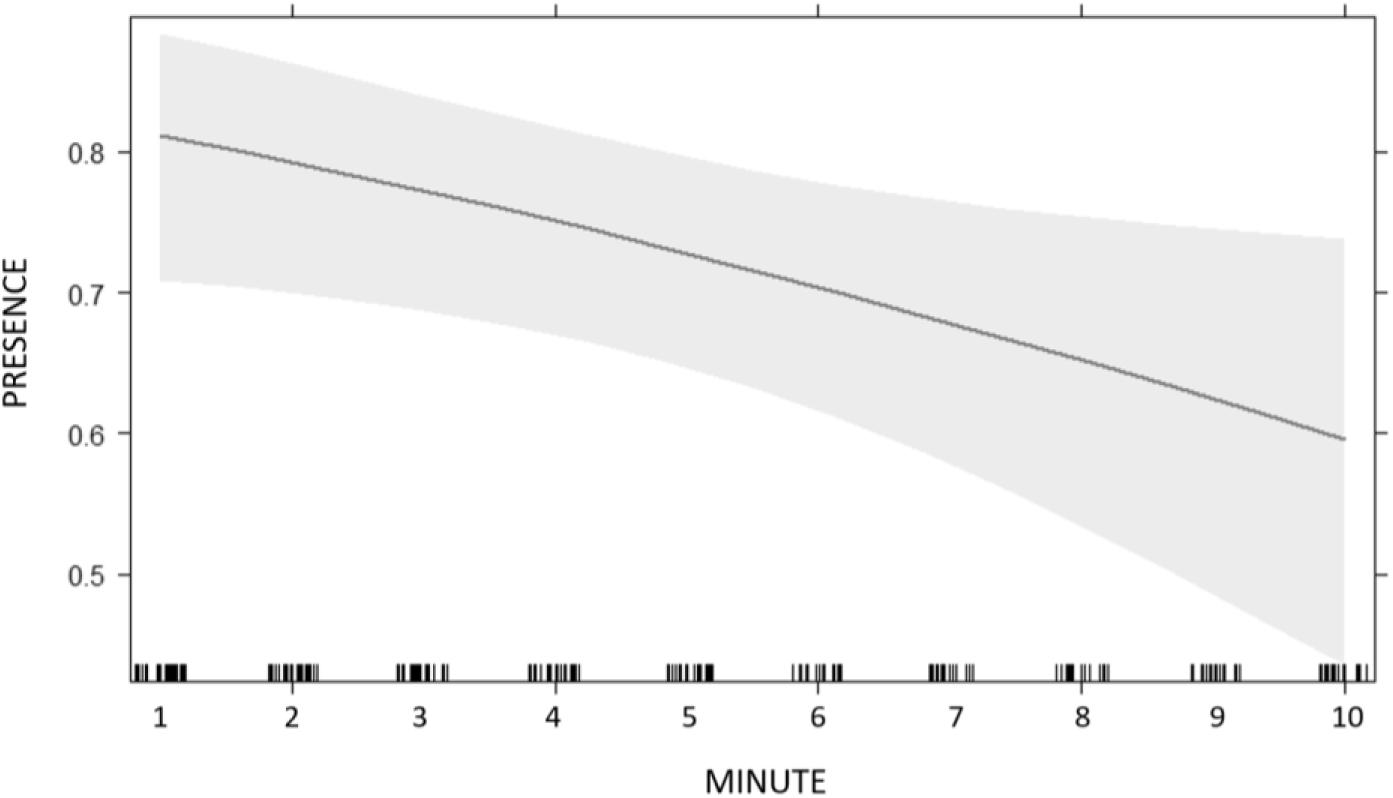
Time variation of anxiety-related behaviours.

**Table 4.** Full results of: GLMM_5_ on factors influencing the presence of anxiety behaviour; GLMM_6_ on factors influencing the presence of anger behaviour; and GLMM_7_ on factors influencing the presence of talking with gestures. For all models the opponent’s identity was included as random factor. The sample size is of N=226 records for all models.

| Predictors | Estimates | SEM | CI<sub>95</sub> | Effect size | $\chi^2$ | p |
| --- | --- | --- | --- | --- | --- | --- |
| **GLMM<sub>5</sub>** | Full vs. null model: $\chi^2=4.417$, df=1, p<0.05 | | | | | |
| (Intercept) | 2.177 | 0.925 | -0.12, 3.33 | 1.61 | 2.354 | **0.019** |
| Minute | -0.119 | 0.054 | -0.66, -0.04 | -0.35 | -2.190 | **0.029** |
| Sex (male) | -0.561 | 0.930 | -2.38, 1.26 | -0.56 | -0.604 | 0.546 |
| Intensity (medium) | -0.999 | 0.455 | -1.89, -0.11 | -1.00 | -2.197 | **0.028** |
| Intensity (high) | 0.414 | 0.432 | -0.43, -1.26 | 0.41 | 0.957 | 0.338 |
| **GLMM<sub>6</sub>** | Full vs. null model: $\chi^2=26.215$, df=1, p<0.001 | | | | | |
| (Intercept) | -1.679 | 1.358 | -5.61, -0.32 | -2.96 | -1.326 | 0.216 |
| Minute | -0.268 | 0.090 | -1.30, -0.27 | -0.78 | -2.997 | **0.003** |
| Sex (male) | 0.695 | 1.398 | -2.04, 3.43 | 0.70 | 0.497 | 0.619 |
| Intensity (medium) | 0.854 | 0.676 | -0.47, 2.18 | 0.85 | 1.263 | 0.206 |
| Intensity (high) | -0.733 | 0.699 | -2.10, 0.64 | -0.73 | -1.048 | 0.295 |
| **GLMM<sub>7</sub>** | Full vs. null model: $\chi^2=6.467$, df=1, p<0.05 | | | | | |
| (Intercept) | -2.061 | 1.215 | -4.27, 0.37 | -1.95 | -1.696 | 0.090 |
| Minute | 0.022 | 0.063 | -0.29, 0.42 | 0.07 | 0.360 | 0.719 |
| Sex (male) | 0.510 | 1.248 | -1.94, 2.96 | 0.51 | 0.408 | 0.638 |
| Intensity (medium) | 1.101 | 0.657 | -0.19, 2.39 | 1.10 | 1.675 | 0.094 |
| Intensity (high) | -0.780 | 0.656 | -2.06, 0.51 | -0.78 | -1.189 | 0.234 |

For GLMM_6_ (target variable: presence/absence anger behaviour) the full model (including all fixed factors: timing, opponents’ sex, /conflict intensity) significantly differed from the null model including the random factor only (likelihood ratio test: χ^2^=16.477, df=4, p<0.01). The only variable that had a significant effect (p<0.05) on the occurrence of anger behaviour was timing (minute) (see full results in Table 4, Figure 5). Conversely, conflict intensity and opponents’ sex had no significant effect (Table 4).

**Figure 5.**
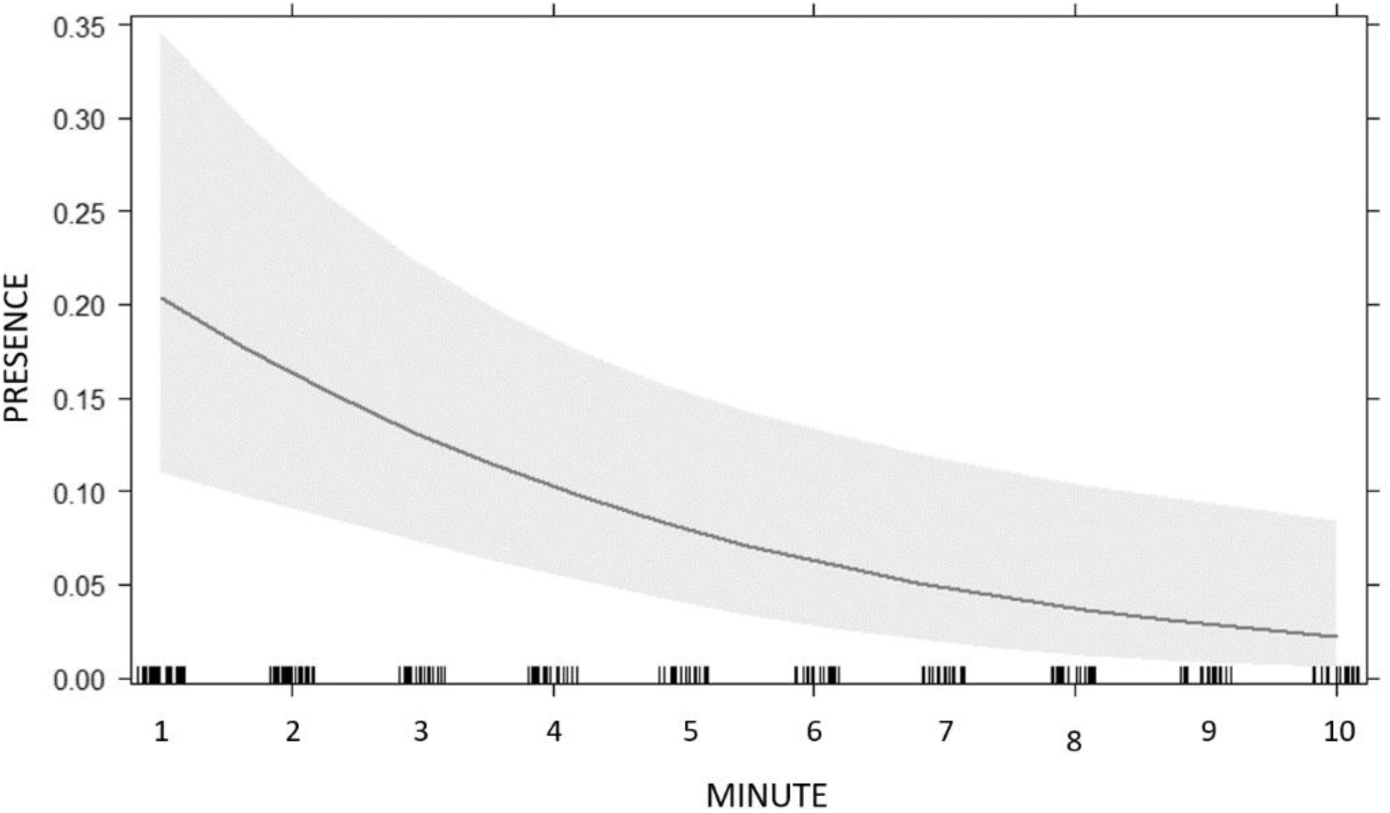
Time variation of anger-related behaviours

For GLMM_7_ we found no significant difference between the full and the null model (likelihood ratio test: χ^2^=7.314, df=4, p=0.121) so none of the predictors had not effect on talking with gestures and we did not proceed with the drop 1 procedure.

## Discussion

Our results show that displacement behaviours (related to anxiety), aggressive behaviours (related to anger) and talking with gestures (possibly related to anxiety/anger), but not body postures, increased in Post-Conflict (PC) compared to a Matched-Control (MC) situation (Prediction 1a and 1b confirmed; Prediction 1c partly confirmed; Table 3, Figure 2). Moreover, both displacement activities and aggressive patterns decreased in the following 10 minutes (Prediction 2a and 2b confirmed; Table 4, Figures 4 and 5), whereas no temporal variation was detected for talking with gestures (Prediction 2c not confirmed; Table 4). Finally we found no effect of conflict intensity for aggressive patterns (Prediction 3b not confirmed) and for talking with gestures (Prediction 3c). Instead, in partial disagreement with our expectation, we found that displacement behaviours were higher after high and low intensity conflict but not after medium intensity conflicts (Prediction 3a not confirmed; Figure 3).

By applying the PC/MC methodology, we were able to show that, compared to a control condition, in the aftermath of a conflict event people displayed social tension behaviours related to a state of anxiety and anger, similarly to what has been reported in other primate species (Aureli 1997; Aureli & van Schaik 1991; Fraser et al., 2008; Palagi & Norscia, 2011; Palagi et al., 2014; Pallante et al., 2018; Romero et al., 2009, 2011). Such behaviours in our case included talking with gestures (increasing after a conflict), but apparently not postures (showing no variation), possibly because the postures included in this studies may also refer to different emotional domains (e.g., deceit, apathy, confidence; Eunson, 2015). In nonhuman primates, aggressive events generate social tension because they can potentially cause direct harm to the opponents and/or jeopardize social stability and therefore the survival of group members (as it occurs in communities where individuals regularly share social relationships; Aureli & de Waal, 2000). The occurrence of a conflict puts the opponents at risk, especially victims, as they are more likely to receive further aggression, which creates a state of uncertainty enhancing opponents’ vigilance and increasing the level of anxiety (Aureli 1997; Aureli & van Schaik, 1991). Hence - beyond the aggression redirected by the previous opponents to uninvolved bystanders (detected in this study) - renewed aggression between opponents shall be taken into account to understand the functions of post-conflict behaviours in humans. Moreover, the level of distress and aggressive manifestations (informed by cortisol and testosterone variations) can increase in case of aggression both with group members or other conspecifics (Cheng et al., 2021; Muller, 2017; Samuni et al., 2019; Wittig et al., 2015). In humans, cortisol and testosterone variations – responsible for distress related anxiety and aggression behaviours – are associated with competition, especially between men (even strangers; Casto & Edwards, 2016). Consistently, regardless of the social relationship, the tension deriving from social challenges are able to activate shared physiological responses (in non-human primates and humans, Preston & de Waal, 2002). We did not take social relationship between opponents into consideration for our analysis of the behaviours of interest. However, the relationship shared between individuals can modulate the physiological response (e.g., testosterone level; Negrey et al., 2023) and the extent to which social tension behaviour increases after a conflict, for example in non-human primates (Aureli, 1997). Because previous research found that in naturalistic observational studies it is possible to infer interpersonal relationships by looking at specific non-verbal cues (Liebst et al., 2023), future studies may use a similar approach and investigate how social bond can affect anxiety and anger expression also in humans.

Our results show that social tension behaviours decrease as the minutes after the conflict elapses. This variation is similar to that observed in nonhuman primates, where the expression of anxiety and anger-related behaviours is higher soon after the end of the conflict, but it returns to baseline levels in the subsequent ten minutes (Aureli & van Schaik 1991; Castles & Whiten, 1998; Cooper et al., 2007; Das et al., 1998; Koski & Sterck, 2007; Kutsukake & Castles, 2001; Pallante et al., 2018). This decrease reflects a drop in the physiological response (following the exposure to a stressor) to restore homeostasis (Maestripieri et al., 1992). Such physiological drop – also present in humans (Herman et al., 2016; Marques et al., 2022) – in nonhuman primates may be accelerated by affiliation between bystanders and opponents (Clay & de Waal, 2013; Fraser et al., 2008; Palagi & Norscia, 2013; Palagi et al., 2014; Pallante et al., 2018; Romero et al., 2010). This aspect could not be explored in this study, but deserves further investigation.

We found that the intensity of the conflict did not influence the occurrence of anger-related behaviours and talking with gestures. This finding seems not to be in line with previous literature reporting that in nonhuman primates (anger-related) aggressive behaviours can be higher in case of severe conflicts (Aureli et al., 1989; Palagi et al., 2008; Palagi & Norscia, 2011; Wittig & Boesch, 2003). This finding may be explained with the observation that the occurrence of anger-related behaviours (after an initial increase) plummets after the conflict (Figure 5), whereas anxiety-related behaviours decrease is delayed (Figure 4). Therefore, the faster decrease of anger-related behaviours might have left less room for the intensity of aggression to modulate such behaviours in the minutes following the conflict.

On the contrary, we found that anxiety behaviours were expressed at higher rates after low and high intensity conflicts compared to intermediate, mild intensity conflicts. This finding may be due to the fact that particularly low intensity aggression generates an unclear situation that can go toward either escalation into more severe fight or calm restoration. This situation can generate moderate anxiety increase, as in humans anxiety can be expressed in case of motivational conflict related to uncertainty (Barker et al., 2018). On the other hand, high intensity aggression may elicit acute stress-related anxiety, as this emotion is also expressed in presence of imminent threats, and fear may activate the distress physiological system (Timmers et al., 2018). One possible reason underlying the lack of variation in correspondence with the intermediate aggression level may be that mild aggression was defined as fights with just one, clear, contact pattern, with no repetition (Table 2). This may make it clearer for the observer when a fight starts and when it is actually over. In general, the way the types of aggression (more or less intense or physical) are categorized is relevant on the likelihood of detecting a modulating effect. For example, in non-human primates, several case studies found no variation in displacement activities between low and high intensity aggression, with the distinction being mostly based on whether physical contact was present in the interaction or not (e.g., Castles & Whiten, 1998; Kutsukake & Castles, 2001). Such categorization may not always be effective – however – as certain types of non-physical aggression may be perceived as severe (e.g., involving screaming and/or chasing; Arnold & Whiten, 2001; Mercier et al., 2019). In humans, the mere presence of a contact may not be sufficient to categorize an aggression as severe. Indeed, a previous study did not consider the mere presence of physical contact as a criteria to discriminate more or less intense conflicts (Lindegaard et al., 2022). As a matter of fact, in humans physical and non-physical aggression may elicit different emotional outcomes in the recipient depending on the type of physical/non-physical aggression (Bernaldo-De-Quirós et al., 2015; Lawrence et al., 2009). However, previous studies were limited by low numbers of conflicts involving high severity aggression, which is also the case in our study that involved 12 of high severity incidences. Future studies should include larger variation in severity levels of the conflicts studied in order to establish how severity matters for emotional displays.

In conclusion, this study documented that the aftermath of a conflict event was characterized by enhanced social tension behaviours in the former opponents, but points toward the possibility that anxiety- and anger-related behaviours do not go in tandem in the post-conflict period. While anger-related behaviours increased regardless of the aggression intensity in a seemingly all-or-nothing manner (as expected in the flight or fight response; McCarty, 2016), the level of anxiety-related behaviours was more sensitive to aggression intensity. Because either low or high aggression intensities could boost anxiety-related behaviours - possibly in relation to mild or acute distress, respectively - future studies may investigate whether more or less severe anxiety is expressed in a different way by individuals in the aftermath of a conflict. Indeed, we showed that social tension behaviours cannot be clumped together because they can be expressed in a fine-grained way, possibly depending on the emotions that they may convey. Despite the limitation of non being able to consider vocalisations (informative of arousal levels) in our analyses - something that we encourage for future investigation - this study was able to highlight the importance of complementing post-conflict research with an ethological approach.

Future studies may investigate how different social tension behaviours are modulated by the post-conflict reunion between former opponents (reconciliation: Aureli, 1997) or affiliation with uninvolved third parties (triadic contacts; Clay & de Waal, 2013; Palagi & Norscia, 2013), as it is known from non-human primates that post-conflict tension behaviours are sensitive to friendly contacts. Our study represents a starting point to define a clearer picture of post-conflict social dynamics and to understand them from a comparative and evolutionary perspective.

## Acknowledgments

We wish to thank Lucia Berti for the drawings. This study was funded by the Dutch Research Council (NWO VI.Vidi.195.083) by means of a grant awarded to Marie Rosenkrantz Lindegaard. This project has also received funding from the European Union’s Horizon 2020 research and innovation program under the Marie Sklodowska-Curie grant agreement No. 101029161 awarded to Virginia Pallante.

## Author contributions

Conceptualization: V.P. and M.R.L. Methodology: V.P., I.N., and M.R.L. Formal Analysis: V.P., I.N. Investigation: V.P., I.N., and M.R.L. Data Curation: V.P. and M.R.L. Writing - Original Draft Preparation: V.P., I.N., and M.R.L. Writing - Review and Editing: V.P., I.N., and M.R.L. Supervision: M.R.L. Project Administration: M.R.L. Funding Acquisition: V.P. and M.R.L. All authors have read and agreed to the published version of the manuscript.

